# Rescue of proteotoxic stress and neurodegeneration by the Zn^2+^ transporter ZIP7

**DOI:** 10.1101/2023.05.22.541645

**Authors:** Xiaoran Guo, Morgan Mutch, Alba Yurani Torres, Maddalena Nano, Drew McDonald, Zijing Chen, Craig Montell, Wei Dai, Denise J. Montell

## Abstract

Proteotoxic stress drives numerous degenerative diseases. In response to misfolded proteins, cells adapt by activating the unfolded protein response (UPR), including endoplasmic reticulum-associated protein degradation (ERAD). However persistent stress triggers apoptosis. Enhancing ERAD is a promising therapeutic approach for protein misfolding diseases. From plants to humans, loss of the Zn^2+^ transporter ZIP7 causes ER stress, however the mechanism is unknown. Here we show that ZIP7 enhances ERAD and that cytosolic Zn^2+^ is limiting for deubiquitination of client proteins by the Rpn11 Zn^2+^ metalloproteinase as they enter the proteasome in Drosophila and human cells. ZIP7 overexpression rescues defective vision caused by misfolded rhodopsin in Drosophila. Thus ZIP7 overexpression may prevent diseases caused by proteotoxic stress, and existing ZIP inhibitors may be effective against proteasome-dependent cancers.

**One-Sentence Summary:** Zn^2+^ transport from the ER to the cytosol promotes deubiquitination and proteasomal degradation of misfolded proteins and prevents blindness in a fly neurodegeneration model.

## Introduction

Protein misfolding and/or aggregation drives numerous diseases including autosomal dominant retinitis pigmentosa, Alzheimer’s, Parkinson’s, Huntington’s and motor neuron diseases, ischemia reperfusion injury, Type I diabetes, and inflammatory bowel disease (*1*–*4*). During normal cellular life, up to 30% of protein molecules misfold and are targeted for degradation by the proteasome (*5, 6*). Some proteins are naturally more prone to misfolding than others and some cells experience greater protein folding challenges than others. Mutations can increase the susceptibility of proteins to misfolding, as can small protein aggregates that seed larger ones. Neurons, and other long-lived cells like stem cells as well as secretory cells are especially dependent on proteasome activity for survival. Unfortunately proteasome activity declines with age (*7, 8*).

Healthy cells degrade misfolded proteins. If the load of misfolded proteins increases, inducing endoplasmic reticulum (ER) stress, cells adapt by activating the unfolded protein response (*9*): expanding the ER, enhancing ER-associated degradation (ERAD), reducing the rate of new protein synthesis, and altering transcriptional programs. If the stress resolves, the cell recovers. If the stress persists and overwhelms the adaptive response, cells die. There is great interest in identifying mechanisms that enhance ERAD in anticipation that such approaches will prevent or reverse degenerative diseases and promote healthy aging (*10, 11*). Conversely, inhibitors of proteasomal degradation of misfolded proteins are in clinical use to treat cancers, especially malignancies of highly secretory cells, such as B cells.

ZIP7 is an evolutionarily conserved, ER Zn^2+^ transporter that promotes cell survival and migration in diverse cell types and organisms (*12*–*15*). ZIP7 is overexpressed in multiple cancers, promotes intestinal stem cell maintenance and is required for B cell differentiation (*12, 16*). Loss of ZIP7 causes ER stress in a variety of cell types. However the mechanism(s) by which ZIP7 mitigates ER stress and contributes to these diverse biological functions are unknown (*17*), and the effects of ZIP7 overexpression are largely unexplored.

We previously identified the *Drosophila* ortholog of ZIP7 (dZIP7, aka Catsup) in a screen for mutations that disrupt border cell migration (*18*) and in a border cell gene expression profile (*19*). Border cells in the *Drosophila* ovary provide an *in vivo* model of collective cell migration that is amenable to unbiased genetic screening (*20*). Here we show that dZIP7 promotes border cell migration by mitigating ER stress. Induction of ER stress by expressing a misfolded protein (Rh1^G69D^) (*21, 22*) blocks border cell migration. Remarkably, overexpression of dZIP7 is sufficient to degrade misfolded Rh1^G69D^, prevent ER stress, and thereby rescue border cell migration. We further show that ER to cytosol Zn^2+^ transport is rate-limiting for ERAD, and specifically for the obligatory deubiquitination of client proteins by the Zn^2+^ metalloproteinase Rpn11. dZIP7 overexpression in photoreceptor cells is sufficient to prevent Rh1^G69D^-induced retinal degeneration. These results illuminate a previously unappreciated rate-limiting requirement for Zn^2+^ in proteasome activation and suggest ZIP7 overexpression as a potential gene therapy for autosomal dominant retinitis pigmentosa and other degenerative diseases.

## Results

### dZIP7 promotes border cell migration and prevents ER stress

*Drosophila* ovaries are composed of ovarioles, which are strings of egg chambers (Fig. 1A) progressing through 14 stages of development, culminating with mature eggs. Each egg chamber is composed of 15 nurse cells and one oocyte (germ cells), surrounded by ∼850 epithelial follicle cells. At stage 9 (Fig. 1B), 4-8 border cells round up at the anterior end of the egg chamber, delaminate from the follicular epithelium, and migrate posteriorly, reaching the anterior border of the oocyte by stage 10. Border cell clusters are composed of 4-6 migratory cells that surround and carry two non-migratory polar cells. Expression of dZIP7 RNAi in the outer, migratory border cells using *fruitlessGal4 (23)* inhibited migration (Fig. 1C, G). The defect was rescued by co-expression of UAS-dZIP7::V5 (Fig. 1D, G). Reduction of dZIP::GFP confirmed the effectiveness of the RNAi (Fig. 1E, E’ and F, F’). Border cell migration was also impaired when dZIP7 RNAi was driven by the *c306Gal*4 (Fig. 1G), which is expressed in both polar and migratory cells. FruitlessGal4-driven RNAi impaired border cell migration at least as much as c306Gal4, indicating that dZIP7 was primarily required in the outer, migratory cells (Fig. 1G). This was further supported by mosaic clone analysis, in which the severity of border cell migration defects were proportional to the number of outer, migratory cells that were homozygous mutant (Supplementary Fig. 1A-C) and mutant cells were mostly excluded from leading positions (Supplementary Fig. 1D-E).

**Fig. 1:**
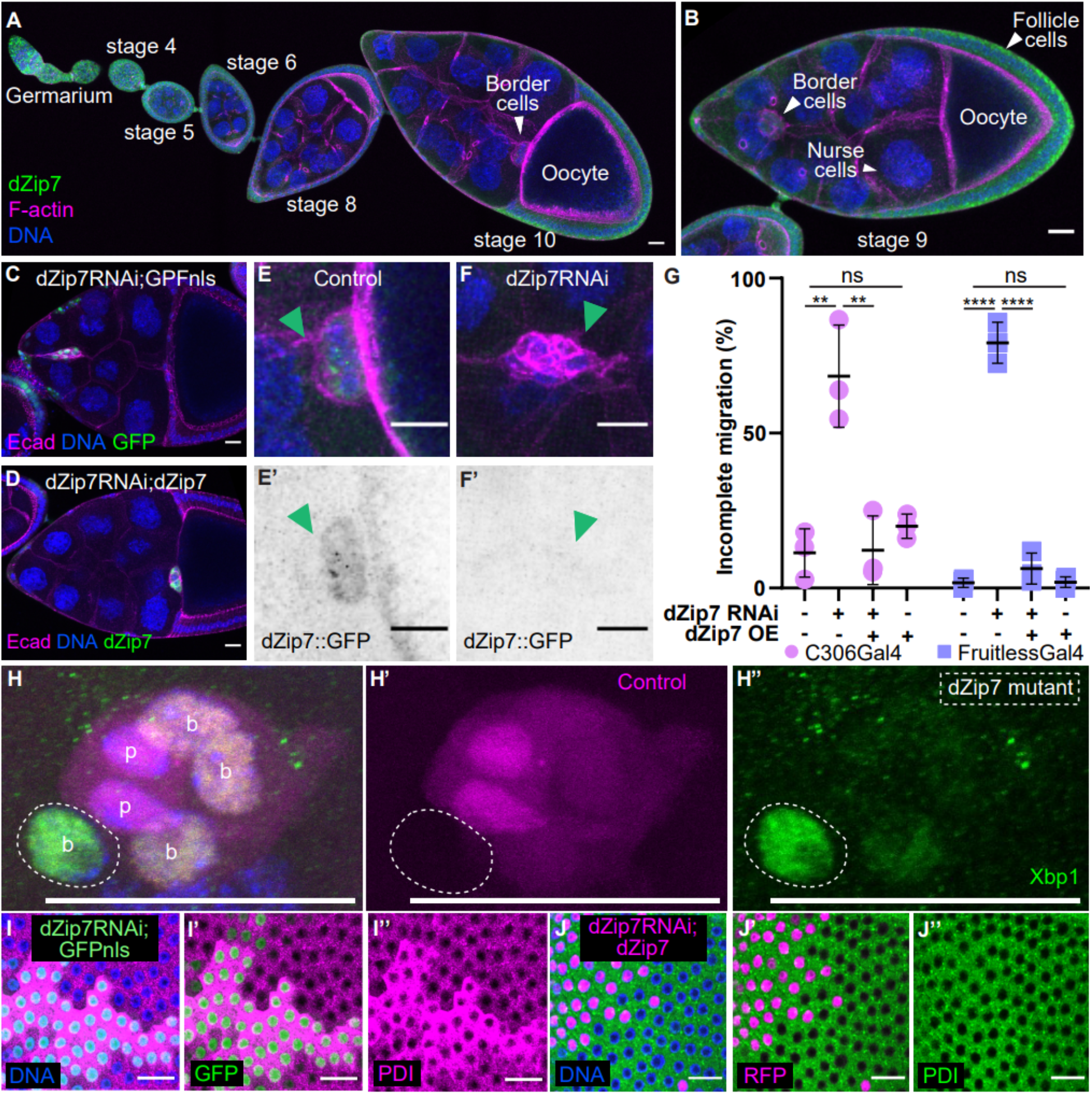
dZIP7 knockdown causes border cell migration defects and ER stress (A, B) Developing Drosophila egg chambers expressing dZIP7::GFP. DNA is in blue. F-actin is in magenta. Border cells migrate during stage 9 (B) and complete migration by stage 10 (A). (C, D) Stage 10 egg chambers with fruitlessGal4 driving expression of UAS-dZiP7RNAi and (C) UAS-GFPnls or (D) UAS-dZIP7::V5 in outer, migratory border cells. (E-F’) dZIP7::GFP expression in control border cells (E, E’) or c306Gal4>dZIP7RNAi (F, F’). (G) Quantification of stage 10 migration defects in c306Gal4 (magenta) and fruitlessGal4 (blue) driving the indicated transgenes. Each dot represents the average of >24 egg chambers (n=3 independent experiments). Error bars=SEM. (H-H”) A mosaic border cell cluster composed of some control cells (RFP+. which can be dZIP7+/+ or dZIP7-/+, magenta, H’) and some homozygous dZIP7 mutant cells (RFP-, outlined). Polar cells (p) express higher levels of RFP compared to outer border cells. Xbp1::EGFP (green. H”) is a marker for ER stress. DNA is in blue. (I-I”) Anti-PDI antibody staining (magenta) reveals that cells expressing dZIP7RNAi and GFPnls exhibit ER expansion. (J-J”) Mosaic clone expressing dZIP7RNAi and dZIP7::V5 and RFP (magenta). Scale bars=20 μm.

dZIP7 is the ortholog of ZIP7, a Zn^2+^ transporter that moves Zn^2+^ from the ER to the cytoplasm and suppresses ER stress in many cell types including mammalian intestinal stem cells (*12, 24*), cancer cells (*24*), Drosophila imaginal discs (*13*) and even in plants (*17*). We confirmed that dZIP7 localizes to the ER in border cells where it co-localized with the ER chaperone PDI much more significantly than with F-actin or the nucleus (Supplementary Fig. 2). Additionally, homozygous mutant dZIP7 border cells expressed the ER stress reporter Xbp1s::EGFP (*25*) whereas dZIP7^+/-^ and dZIP7^+/+^ cells did not (Fig. 1H-H”). We also observed increased expression of the ER chaperone PDI in homozygous dZIP7 RNAi-expressing follicle cell clones (Fig. 1I-I”), a phenotype that we rescued with a wild type, V5-tagged dZIP7 transgene (Fig. 1J-J”). Accumulation of XBP1 and PDI are indicative of induction of an adaptive unfolded protein response (UPR)(*26*). We conclude that cells lacking dZIP7 experience ER stress and impaired migration, raising the question as to whether ER stress inhibits motility.

### dZIP7 is limiting for ERAD

To address whether ER stress inhibits migration, we expressed a misfolded rhodopsin protein, Rh1^G69D^, known to induce ER stress (*27, 28*). Misfolded rhodopsin accumulates in the ER and causes retinal degeneration in flies and humans (*28*–*31*). We found that Rh1^G69D^ accumulated to high levels intracellularly in border cells, induced ER stress, and blocked their migration (Fig. 2A-D), demonstrating that ER stress is sufficient to impair motility.

**Fig. 2:**
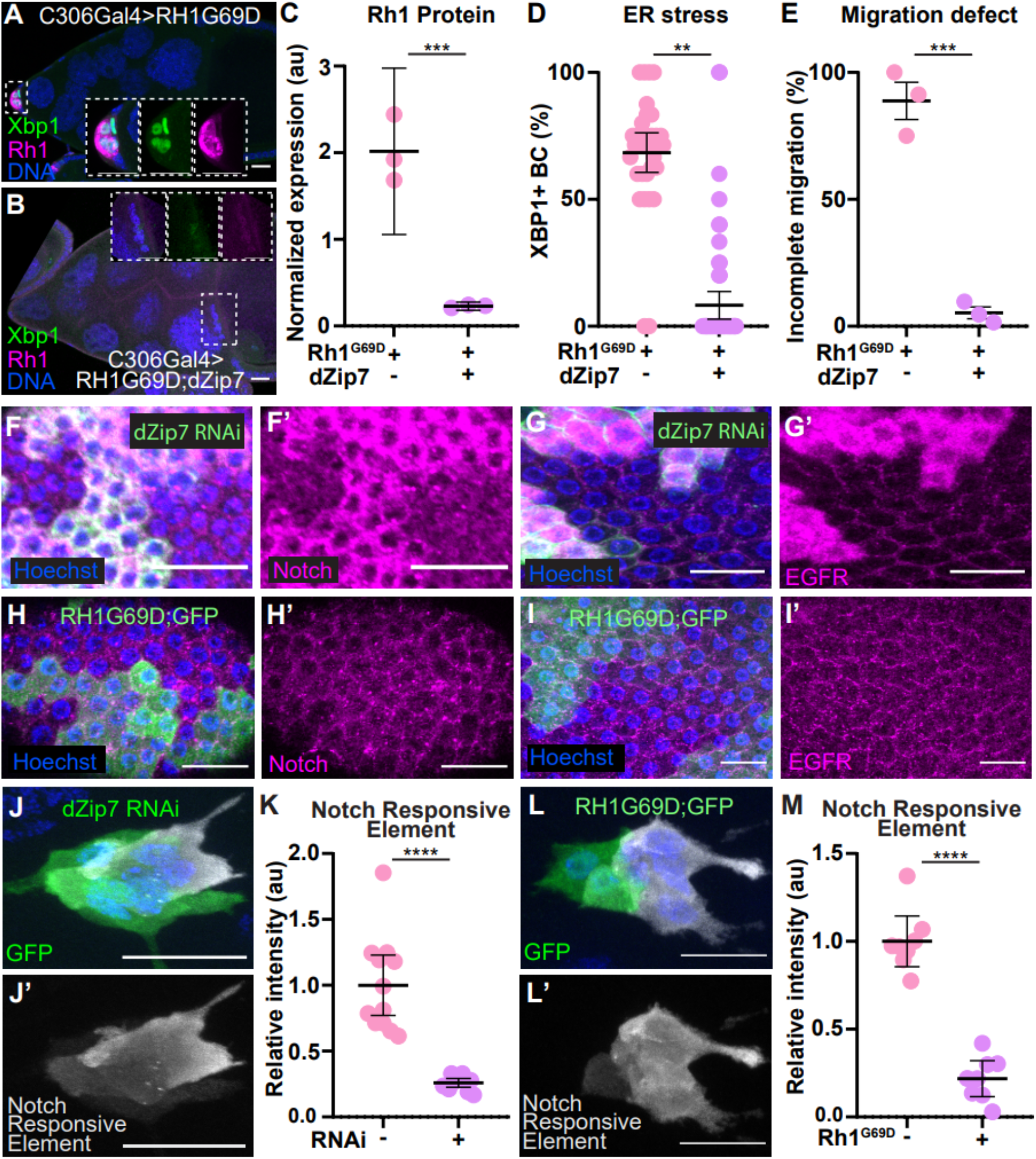
dZIP7-overexpression is sufficient to degrade misfolded Rh1G69D, prevent ER stress, and rescue migration. (A) Stage 10 egg chamber expressing Rh1G69D (magenta) and the ER stress sensor Xbp1::EGFP (green) in border cells using c306Gal4. Rh1G69D accumulation induced ER stress and blocked migration. (B-D) Co-expressing dZip7 reduced Rh1G69D (B-C) and Xbp1::EGFP (B-D) and rescued migration (B-E). (C) Each dot represents one experiment (n>17 clusters). Error bars=95% confidence intervals. (D) Each dot represents an individual border cell. Error bars=95% confidence intervals. (E) Each dot is the average of all egg chambers in one experiment. Error bars=SEM. (F-G’) Intracellular Notch (F, F’) and EGFR (G, G’) accumulated in epithelial follicle cell clones expressing dZIP7RNAi (GFP+, green) relative to neighboring wild type cells. (H-l) Mosaic clones of follicle cells expressing Rh1G69D and GFP. Rh1G69D expression does not cause accumulation of Notch (H, H’) or EGFR (I, I’) relative to wild type cells. (J-M) Notch transcriptional activity visualized with a Notch responsive element reporter (white). (J-K) Notch in dZIP7RNAi-expressing cells (GFP+) compared to neighboring wild type (GFP-) cells. (L-M) Notch in Rh1G69D-expressing cells (GFP+) compared to wild type cells (GFP-). ** P≤0.01, *** P≤0.001. Scale bars=20 μm.

Since loss of dZIP7 caused ER stress, we wondered if dZIP7 overexpression might suppress ER stress, so we co-expressed Rh1^G69D^ and dZIP7::V5 in border cells. Interestingly, dZIP7::V5 expression virtually eliminated Rh1^G69D^ protein (Fig. 2B, C), reduced ER stress (Fig. 2B, D) and restored normal border cell migration (Fig. 2B, E). Together, these results suggest that dZIP7 is a limiting factor for degrading misfolded Rh1, i.e. for ERAD, and its overexpression is sufficient to mitigate the negative biological consequences of ER stress.

Notch and EGFR accumulate abnormally in the ER in ZIP7-deficient fly wing disc cells (*13*) and human cancer cells (*32*). We also observed abnormal accumulation of Notch (Fig. 2F, F’ and Supplementary Fig. 3A, A’) and EGFR (Fig. 2G, G’ and Supplementary Fig. 3B, B’) but not E-cadherin (Supplementary Fig. 3C, C’) in dZIP7 knockdown follicle cell clones, supporting the generality of the phenomenon. Although Rh1^G69D^ expression in follicle cells including border cells caused ER stress and induced the adaptive UPR, neither Notch (Fig. 2H, H’) nor EGFR (Fig. 2I, I’) accumulated abnormally in Rh1^G69D^-expressing cells. Yet, both ZIP7 knockdown cells (Fig. 2J, J’) and Rh1^G69D^-expressing cells (Fig. 2K, K’) exhibited reduced Notch transcriptional responses (Fig. 2L). We conclude that dZIP7 knockdown causes two independent effects on Notch: accumulation of misfolded protein in the ER due to reduced ERAD and inhibition of Notch transcriptional activity, presumably as a consequence of the ER stress response (*33*–*35*).

### dZIP7 promotes Rpn11-mediated deubiquitination of misfolded proteins prior to proteasomal degradation

To investigate which step of the ERAD process requires dZIP7, we used an antibody that recognizes ubiquitinated proteins to compare wild type, dZIP7-overexpressing, and dZIP7 RNAi cells. One hypothesis was that dZIP7 might provide Zn^2+^ to ubiquitin ligases such as Hrd1 and SORDD1/2, which are Zn^2+^-binding proteins that reside in the ER membrane (Fig. 3A). If this were true, we might expect dZIP7 overexpression to increase - and dZIP7 RNAi to decrease - the abundance of polyubiquitinated proteins (PUBs). Surprisingly, we observed the opposite effect. dZIP7 RNAi increased polyubiquitinated proteins (Fig. 3B, B’) compared to the lacZ control (Fig. 3C, C’ and Supplementary Fig. 4A-B’). Conversely, dZIP7 overexpression essentially eliminated detectable polyubiquitinated proteins (Fig. 3D-E and Supplementary Fig. 4C, C’). We also found that Rh1^G69D^ expression increased the accumulation of polyubiquitinated proteins compared to the control (Supplementary Fig. 4D-J’), and ZIP7 overexpression suppressed the effect (Supplementary Fig. 4F, F’, G and J, J’). Since polyubiquitinated proteins accumulated in the absence of dZIP7, we conclude that ZIP7 functions downstream of ubiquitination to promote degradation of misfolded proteins.

**Fig. 3:**
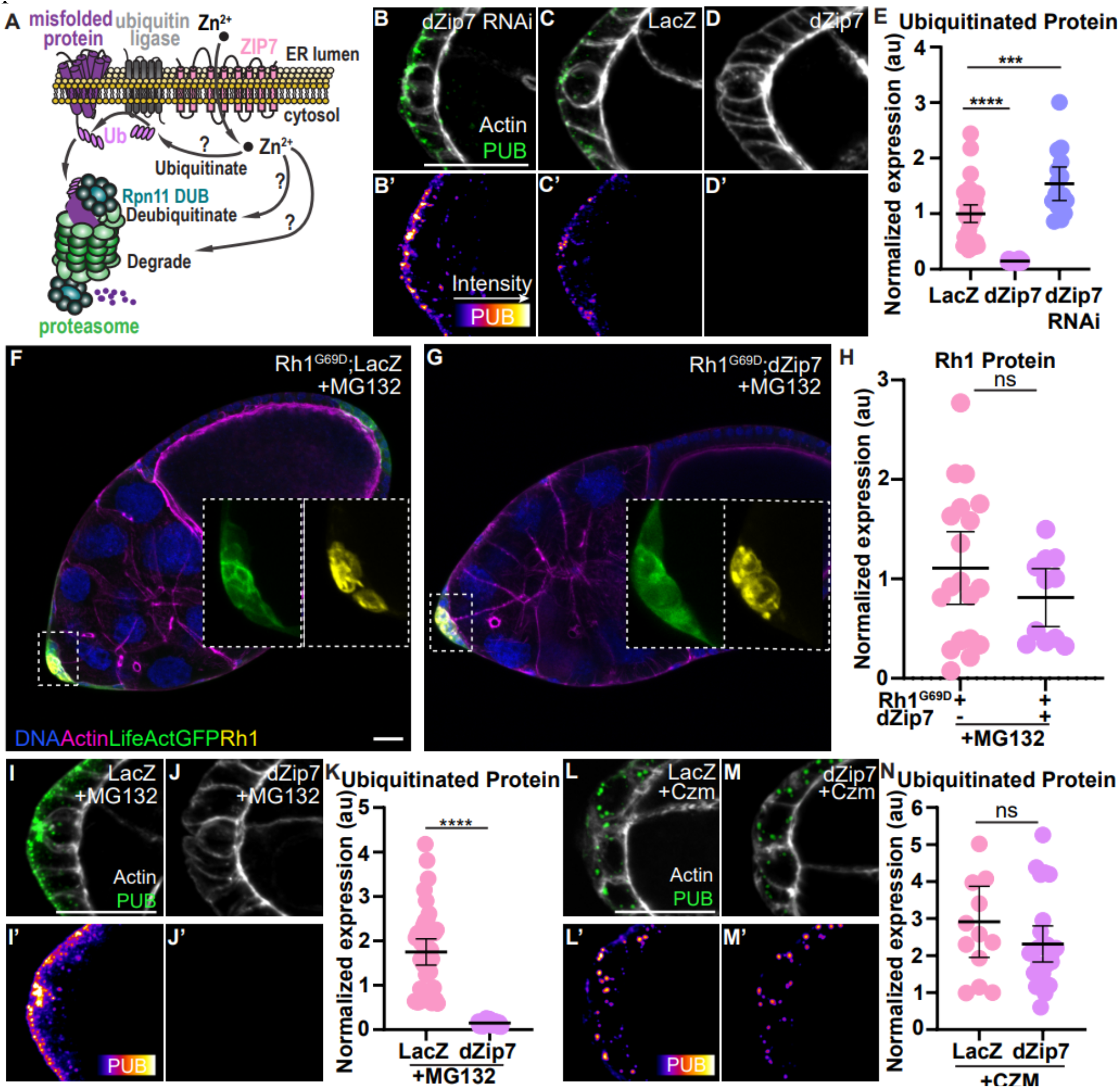
dZIP7 promotes Rpn11-mediated deubiquitination of proteins required for proteasome entry. (A) Schematic of proteasomal processing of misfolded proteins. (B-D’) Representative images of stage 8 egg chambers stained with an antibody against ubiquitinated proteins (PUB. green in B-D. FIRE LUT in B’-D’). (E) Quantification of fluorescence intensity of PUB staining in control (lacZ), dZIP7 overexpression and dZiP7RNAi expressing border cells. (F-H) Stage 10 egg chambers expressing misfolded Rh1G69D in border cells (insets) treated with 10μM of the MG 132 proteasome inhibitor for 5 hours and stained with an antibody against Rh1. dZip7 co-expression with Rh1G69D did not reduce Rh1 protein levels in the presence of MG132 showing that the dZiP7-mediated degradation of Rh1G69D (see Fig. 2B) is mediated by the proteasome and that dZiP7 functions upstream of the 20S core enzyme blocked by MG132. (I-J’) dZip7 overexpression reduced ubiquitinated protein levels in egg chambers treated with MG 132 (10pM, 5 hours) Relative fluorescence intensity is quantified in (K). (L-M’) dZip7 overexpression does not prevent ubiquitinated protein buildup in egg chambers treated with Rpn11 inhibitor Capzimin (Czm, 20μM, 5 hours); this effect is quantified in (N). Dots represent individual border cell clusters. Error bars=95% confidence intervals. *P≤0.05, ** P≤0.01, **** P≤0.0001. Scale bars=20 μm.

The observation that expression of dZIP7 reduced the abundance of polyubiquitinated proteins suggested that it might be required instead for deubiquitination. There are many deubiquitinating enzymes (DUBs), but Rpn11 stood out as a top candidate. Whereas deubiquitination by some DUBs can rescue proteins from degradation, deubiquitination by Rpn11 is an essential prerequisite for entry of client proteins into the proteasome core and thus, like dZIP7, is essential for misfolded protein degradation (*36*) (Fig. 3A). Furthermore, unlike most DUBs, Rpn11 requires Zn^2+^ for catalysis (*37*–*39*). *In vitro*, in the absence of Zn^2+^, ubiquitinated substrates and the 26S proteasome including Rpn11 assemble into inactive complexes and the DUB is activated by addition of Zn^2+ (*37*)^.

Since dZIP7 transports Zn^2+^ to the cytosol (*24*) and promotes loss of ubiquitinated proteins (Fig. 3E) and degradation of Rh1^G69D^ (Fig. 2B, C), we hypothesized that dZIP7 might be limiting for Rpn11 activity. A prediction of this model is that the proteasome inhibitor MG132 would block the ability of dZIP7 overexpression to promote Rh1^G69D^ degradation. In the absence of MG132, ZIP7 overexpression virtually eliminated Rh1^G69D^ protein (Fig. 2B, C). In contrast, MG132 largely prevented the ability of ZIP7 to promote degradation of Rh1^G69D^ (Fig. 3F-H).

MG132 also prevented dZIP7-mediated rescue of the Rh1G69D border cell migration defect (Fig. 3F, G compared to Fig. 2A, B).

MG132 blocks a protease within the core of the 20S proteasome, not Rpn11 itself, so the model further predicts that even in the presence of MG132, ZIP7 should still promote deubiquitination, which we observed (Fig. 3I-K, Supplementary Fig. 4K-L’). We conclude that ZIP7 enhances deubiquitination of misfolded proteins upstream of 20S proteasomal degradation.

If ZIP7 promotes Rpn11-mediated deubiquitination, an Rpn11 inhibitor should block the ability of dZIP7 to enhance deubiquitination of misfolded proteins. To test this prediction, we used the Rpn11 inhibitor capzimin, which specifically binds and blocks the binding of Zn^2+^ within the catalytic site of Rpn11 (*40*). Capzimin largely prevented the effects of ZIP7 on deubiquitination (Fig 3L-N, Supplementary Fig. 4M-N’). We conclude that dZIP7 is limiting for the obligatory Zn^2+^- and Rpn11-dependent deubiquitination of misfolded proteins prior to proteasomal degradation.

### *ZIP7-mediated Zn*^*2+*^ *transport is limiting for ERAD* in Drosophila

ZIP7 resides in the ER membrane and transports Zn^2+^ from the ER to the cytosol (*24, 41*). To test whether the Zn^2+^ transporter activity of dZIP7 is important for border cell migration, we introduced point mutations, H315A and H344A, which replace histidine residues that are required for Zn^2+^ transport (*42*) (Fig. 4A, purple) and are conserved between dZIP7, ZIP7 and a more distant family member from *Arabidopsis* IRT1 (Supplementary Fig. 5). As controls, we engineered dZIP7^H187A^ and dZIP7^H183A^ mutants (Fig. 4A, green), which do not affect Zn^2+^ transport in IRT1 (*43*). We generated transgenic flies expressing the mutants under Gal4/UAS control and included a V5 tag so that we could monitor protein abundance and localization. We then co-expressed each of these RNAi-resistant transgenes with dZIP7 RNAi and evaluated protein expression and border cell migration. The point mutations predicted not to disrupt Zn^2+^ transport, dZIP7^H187A^ and dZIP7^H183A^, rescued border cell migration to nearly wild type levels (Fig. 4B, C) whereas neither dZIP7^H344A^ nor dZIP7^H315A^ provided significant rescue (Fig. 4D, E), as quantified in Fig. 4F. All the proteins were stably expressed and correctly localized to the ER (Fig. 4B-E), therefore the lack of rescue was likely a consequence of impaired transporter activity rather than impaired expression or localization. Similarly, the Zn^2+^-transport-deficient proteins (dZIP7^H344A^ and dZIP7^H315A^) failed to rescue ER accumulation of Notch and EGFR, when clonally co-expressed with dZIP7 RNAi in a subset of border cells. Notch and EGFR accumulated abnormally in the cells co-expressing dZIP7^H344A^ or dZIP7^H315A^ with dZIP7 RNAi (Fig. 4G-J’, magenta nuclei) compared to neighboring control cells (blue nuclei). In contrast, dZIP7 RNAi-expressing cells that co-expressed dZIP7^H187A^ or dZIP7^H183A^ cells (magenta nuclei in Fig. 4K-N’) exhibited similar Notch and EGFR levels as the neighboring control cells (blue nuclei in Fig. 4K-N’). The results are quantified in Fig. 4O,P. From these experiments, we conclude that Zn^2+^ transport is an essential function of dZIP7 in promoting ERAD.

**Fig. 4:**
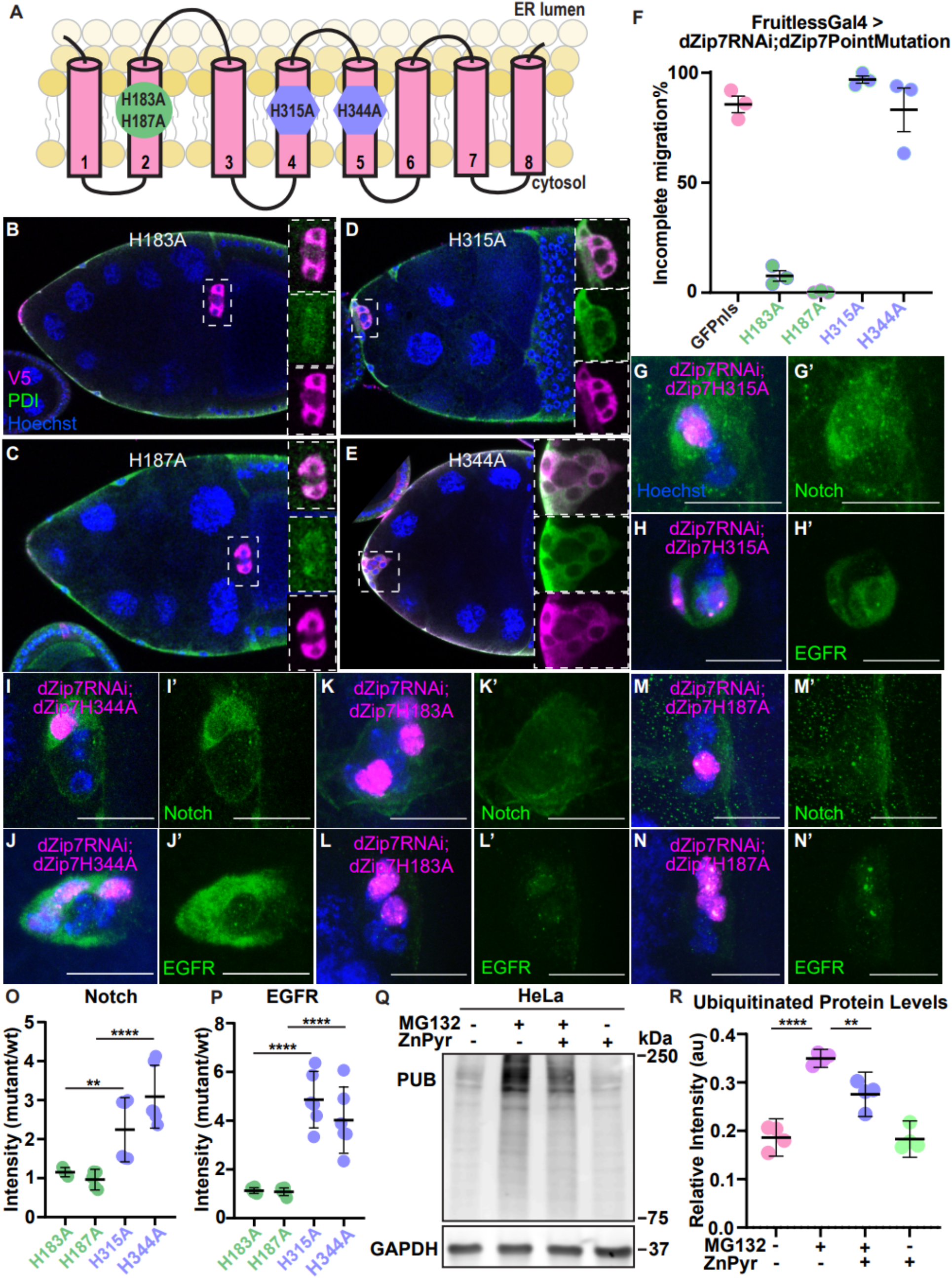
Cytosolic Zn2+ is limiting for ERAD. (A) Schematic of transmembrane domains and topology of dZIP7. Point mutations H183A and H187A reside within the second transmembrane domain while H315A and H344A are within the highly conserved HELP domain and CHEXPHEXGD motif on the fourth and fifth transmembrane domains required for Zn2+ transport. (B-E) Co-localization of V5-tagged, RNAi-resistant dZIP7 mutants with the ER marker PDI (green) in border cells. (F) Quantification of incomplete migration at stage 10 in egg chambers expressing dZIP7RNAi with the indicated mutant forms. N=3 independent experiments. Error bars=SEM. (G-N’) Mosaic expression of dZIP7RNAi together with the indicated mutant forms of dZIP7 marked by RFPnls (magenta) and stained for Notch or EGFR (green). Scale bars=20 μm. Quantification of the fold change of Notch (O) and EGFR (P) expression in dZip7 mutants. (Q) Representative western blot on HeLa cell protein extract probed for polyubiquitinated protein and GAPDH. Cells were treated with/without proteasome inhibitor MG132 and zinc pyrithione (ZnPyr), a zinc ionophore. Treatments from left to right: DMSO, 500nM MG132, 500nM MG132 + 1μM ZnPyr, 500nM ZnPyr. (R) Adding ZnPyr to MG132-treated cells reduces polyubiquitinated protein levels. Error bars=95% confidence intervals. ^**^P≤0.01, ^****^P P≤0.0001.

### Zn^2+^ is limiting for deubiquitination of proteasome client proteins in human cells

ZIP7 is nearly ubiquitously expressed in cells from organisms as diverse as plants and animals (*17*). Our results, combined with the observations that the proteasome is highly abundant (*36*) whereas cytosolic [Zn^2+^]_free_ is extremely low (∼1 nM) (*24*), suggested that Zn^2+^ might be rate limiting for proteasome activity. So, we tested the ability of a Zn^2+^ ionophore, pyrithione (an organic salt of zinc capable of permeating cell membranes) (*24*) to enhance deubiquitination of proteins in human cells.

This ionophore has previously been shown to rescue ER stress in ZIP7-deficient HeLa cells (*24*). So, we incubated HeLa cells with MG132 to induce ER stress and increase the abundance of polyubiquitinated proteins (Fig.4Q). We then tested the effect of the Zn^2+^ ionophore. The added Zn^2+^ reduced the accumulation of polyubiquitinated proteins (Fig 4Q). We conclude that cytosolic Zn^2+^ is limiting for proteasomal degradation of misfolded proteins.

### dZIP7 overexpression prevents retinal degeneration caused by Rh1^G69D^

The observations that dZIP7 overexpression is sufficient to degrade Rh1^G69D^, reduce ER stress, and rescue border cell migration and the ubiquity of ZIP7 and proteasomes suggested that dZIP7 overexpression might also be effective at suppressing retinal degeneration due to folding-defective rhodopsin. To test this hypothesis, we co-expressed UAS-dZIP7::V5 with UAS-Rh1^G69D^ in fly photoreceptor cells using GMR-Gal4. Eye morphology was normal in flies expressing dZIP7 alone (Fig. 5A, B and G) whereas Rh1^G69D^ causes severe disruption of eye morphology compared to controls (*21, 27*–*29, 31*) and (Fig 5C, D and G). Co-expression of Rh1^G69D^ and dZIP7 fully rescued eye morphology in the majority of flies examined (Fig. 5E, F and G).

**Fig. 5:**
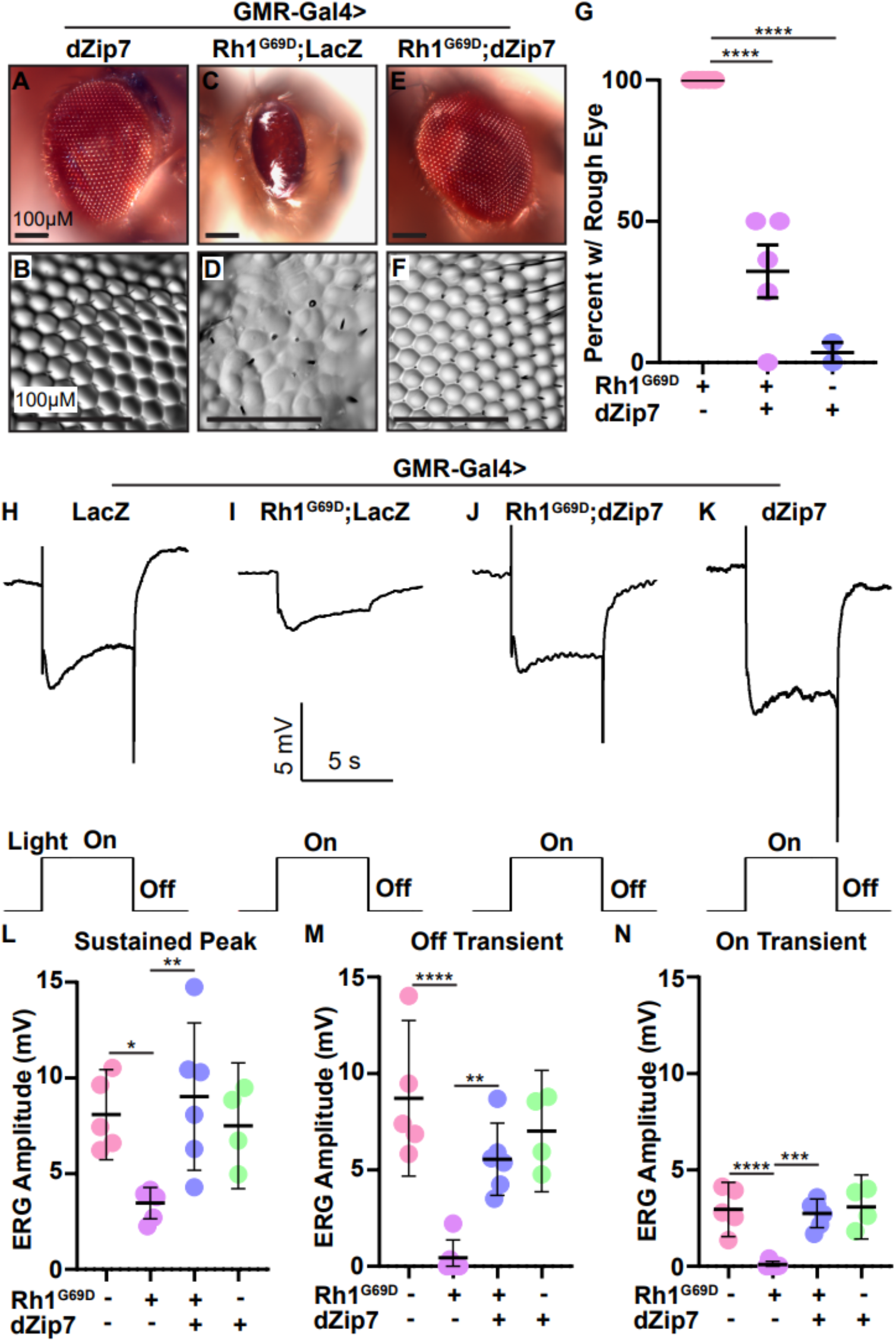
dZIP7 overexpression prevents Rh1G69D retinal degeneration. (A, C, E) Representative light photomicrographs of retinal morphology. (B, D, F) DIC images of retinal imprint morphology. (G) Quantification of number of flies with rough eye. dZip7 overexpression reduces the proportion of flies with rough eye morphology. Each dot represents an average of >10 flies observed in each experiment. (H-K) Representative ERG recordings of one-week-old flies. (L-N) Quantification of ERG recordings. dZip7 co-expression with Rh1G69D returns ERG amplitude to control levels. Error bars represent 95% confidence intervals. *P≤0.05, ** P≤0.01, **** P≤0.0001.

To test whether dZIP7 overexpression could restore visual function, we carried out electroretinogram (ERG) recordings, which measures the summed responses of all retinal cells to light. Control flies display a corneal negative receptor potential upon turning on a light stimulus, which quickly decays to baseline upon termination of the light (Fig. 5H). The large maintained component of the ERG results principally from activation of the phototransduction cascade. The on- and off-transient responses, which are nearly coincident with the initiation and cessation of the light stimulus (Fig. 5H), depends on synaptic transmission from the photoreceptor cells to postsynaptic cells in the optic lobes. Expression of Rh1^G69D^ in photoreceptor cells greatly diminished the amplitude of the ERG, and eliminated the on- and off-transients (Fig. 5I, L-N). Of note, overexpression of dZIP7 in Rh1^G69D^ photoreceptor cells restored a normal ERG, including a full receptor potential and synaptic transmission, as evidenced by the on- and off-transients (Fig. 5J, L-N). As a control, we found that expression of dZIP7 alone in photoreceptor cells had no adverse effects on the ERG (Fig. 5K-N). These data demonstrate that over-expression of dZIP7 rescued the deleterious impact of Rh1^G69D^ on the response of retinal cells to light without side effects.

## Discussion

### A model for dZIP7 function: Zn^2+^ transport from the ER to the cytosol is limiting for ERAD and mitigation of ER stress

dZIP7 is a conserved protein that goes by names including ZRT1 in yeast, IRT1 in plants, dZIP7 or *Catsup* in Drosophila, and SLC39a7/Zip7/Ke4 in mammals. While many studies come to a common conclusion that loss or inhibition of ZIP7 disrupts ER homeostasis in cells from plants to flies and humans (*12, 17, 24, 32, 44, 45*), the mechanism has been unclear (*17*). The data presented here provide evidence for an unanticipated mechanism. Our data support the model that dZIP7 promotes ERAD and prevents ER stress by providing free Zn^2+^ to the catalytic site of the Rpn11 DUB in the proteasome lid (Fig. 6A). This role for dZIP7 is critical since free Zn^2+^ is present at exceedingly low intracellular levels (*24*), and is therefore limiting. In the absence of ZIP7, misfolded/unfolded proteins accumulate and cause ER stress (Fig. 6B and C).

**Fig. 6.**
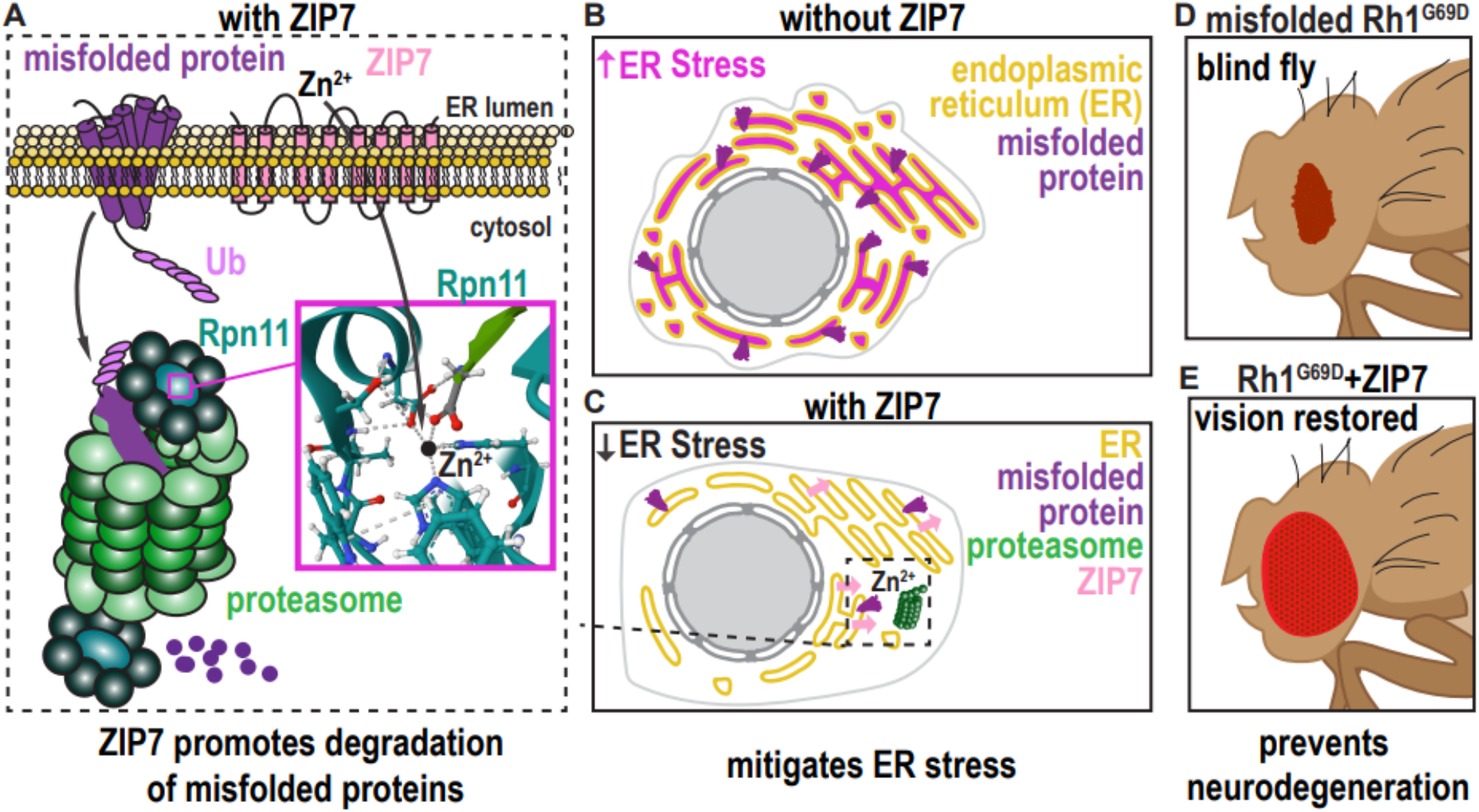
Model for ZIP7 function at molecular, cellular, and organismal scales A) ZIP7 transports Zn2+ from ER lumen to the cytosol where it is limiting for the activity of the Rpn11 deubiquitinase in the lid of the proteasome. B) When ZIP7 is knocked down or inhibited, misfolded proteins accumulate in the ER and cause ER stress and the UPR. C) In the presence of ZIP7 misfolded proteins are degraded and ER stress is prevented D) In a fly model of autosomal dominant retinitis pigmentosa, the Rh1G69D mutant form of rhodopsin causes ER stress, photoreceptor cell death and blindness. E) Overexpression of ZIP7 prevents ER stress, cell death, and blindness.

Furthermore, we show that dZIP7 overexpression is sufficient to enhance proteasomal degradation of misfolded proteins, including Rh1^G69D^, preventing the harmful effects of ER stress including blindness (Fig. 6D and E).

In contrast to earlier work that suggested that ZIP7 primarily promotes trafficking of membrane proteins such as Notch and EGFR (*13, 32*), our results show that release of Zn^2+^ from the ER to the cytosol via ZIP7 is limiting for ERAD. We propose that the step in ERAD that is most sensitive to [Zn^2+^] is deubiquitination of client proteins by Rpn11, which is a Zn^2+^ metalloproteinase (*37, 38*). Our *in vivo* genetic and pharmacological studies are supported by *in vitro* biochemistry. Rpn11 requires Zn^2+^ to deubiquitinate client proteins (*37*–*40*). This is an essential step so that the client protein can insert into the 20S proteasome for degradation by trypsin, chymotrypsin, and caspase-like endoproteases. Rpn11 also enhances proteasomal degradation by allowing ubiquitin to be recycled. Worden et al (*37*) were able to assemble a complex *in vitro* composed of a ubiquitinated substrate (insulin) and the 26S proteasome, including Rpn11. In the absence of free Zn^2+^ the complex assembles and is stable but Rpn11 is catalytically inactive, so ubiquitinated substrate accumulates. Upon addition of Zn^2+^, Rpn11 deubiquitinates the client protein. In the presence of the proteasome inhibitor MG132, Rpn11 deubiquitinates the client but it is not degraded, so deubiquitinated protein accumulates.

We observe remarkably similar effects by manipulating dZIP7 *in vivo* as Worden et al observed by manipulating Zn^2+^ *in vitro* (*37*). In the absence of dZIP7, ubiquitinated proteins accumulated, whereas upon overexpression of dZIP7 in the presence of MG132, deubiquitinated substrate proteins (e.g. Rh1^G69D^) accumulate. It is reasonable to propose that cytosolic free Zn^2+^ would be rate limiting for proteasome function because proteasomes are abundant, whereas cytosolic free Zn^2+^ is vanishingly rare at ∼1 nM (*24*), which is ∼100-fold less than the typical free cytosolic [Ca^2+^]. We propose that ZIP7 provides rate-limiting Zn^2+^ to Rpn11, and thus that the level of ZIP7 determines a cells’ capacity to degrade misfolded proteins. In support of this idea and the generality of the mechanism proposed here, we found that increasing intracellular Zn^2+^ enhanced deubiquitination of proteins in a human cell line.

Free Zn^2+^ is exceptionally rare in the cytosol despite the fact that Zn^2+^ is the second most abundant divalent cation in cells because nearly all Zn^2+^ is bound to proteins. While an essential trace element, excess cytosolic Zn^2+^ can be toxic (*46*), yet ZIP7 overexpression did not cause detectable harm either to follicle cells in the ovary or to photoreceptor cells in the eye. This suggests that the Zn^2+^ transported to the cytosol via ZIP7 might predominantly exert its effects locally. ZIP7 may not directly bind to the ERAD machinery though because it was not detected in an extensive proteomic analysis (*47*). The human genome encodes 24 Zn^2+^ transporters, 14 of which belong to the ZIP family which move Zn^2+^ into the cytoplasm from outside the cell or from inside an organelle while 10 are members of the ZnT family which transport Zn^2+^ out of the cell or into organelles (*48, 49*). The large sizes of these families are consistent with the idea that local Zn^2+^ sources may be important for promoting necessary Zn^2+^-dependent processes without increasing global levels, which would be toxic.

The ability of dZIP7 overexpression to alleviate the ER stress and cellular defects due to Rh1^G69D^ expression has some general biomedical implications. Dominant mutations in rhodopsin that impair folding and cause accumulation in the ER cause retinal degeneration in human patients (*30*), for which there is no effective prevention or therapy. Over-expression of proteins that enhance ERAD is a promising therapeutic strategy. Additionally, toxic protein aggregates have been proposed to kill neurons by inhibiting ERAD in numerous neurodegenerative diseases including Huntington’s, Altzheimer’s, Parkinson’s, frontotemporal dementia, and others, even when the toxic protein is not localized in the ER (*2, 50*). Thus, strategies to enhance ERAD may be useful in treating other diseases as well.

The suppression of ER stress and border cell migration by dZIP7 overexpression is consistent with the observation that ZIP7 is over-expressed in numerous cancers where it promotes survival, proliferation and migration and correlates with disease progression, invasion, and metastasis (*14*–*16, 51*). A ZIP7 inhibitor was identified in a screen for drugs to treat Notch-dependent cancers, based on the model that ZIP7 is important for Notch trafficking (*32*). We show that ER stress impairs Notch transcriptional activity independent of any trafficking defect because Rh1^G69D^ inhibits Notch signaling without abnormal Notch or EGFR protein accumulation in the ER lumen. How ER stress or the UPR inhibits Notch signaling is not clear, but the observation that a pharmacological inhibitor of ZIP7 was identified as a suppressor of Notch signaling by the Notch intracellular domain (NICD) in cultured U2OS osteosarcoma cells (*32*) suggests that there is a deeply conserved requirement for ZIP7 for Notch transcriptional activity. Nolin et al (*32*) showed that ZIP7 inhibition causes accumulation of full length Notch and a decrease in the NICD, and concluded that Notch activation by proteolysis was likely perturbed upon inhibition of ZIP7. An alternative interpretation is that full-length Notch accumulates in the ER lumen due to inhibition of ERAD, and that the NICD is independently degraded more rapidly as part of the global ER stress response.

Our results suggest that ZIP7 inhibitors might be effective against cancers that rely especially heavily on proteasomes. Proteasome inhibitors such as bortezomib are approved for the treatment of B cell malignancies including multiple myeloma and mantle cell lymphoma (*52*). Our results suggest that ZIP7 inhibitors might be repurposed to treat those cancers as well, especially considering that resistance typically develops against a single therapeutic agent. Interestingly, hypomorphic mutations in ZIP7 cause a B cell deficiency due to defects in B cell differentiation in human patients (*53*). Although the mechanism underlying this phenotype is unknown, our results implicate ZIP7 in the UPR, and mutations that compromise the UPR also cause B cell deficiency due to defective B cell differentiation. So, the results presented here suggest a possible link between these otherwise disparate observations. B cell development appears to depend upon a functional UPR and ER stress response, perhaps to ensure their resilience to the natural ER stress B cells experience when they secrete large quantities of antibody.

Finally, the similarities in dZIP7 functions and phenotypes across disparate cells, tissues, and organisms suggests that the border cell system offers an excellent model for deciphering the fundamental and conserved effects of this protein *in vivo*.

## Acknowledgments

Thanks to Dr. Diego Acosta-Alvear for sharing his expertise in ER stress responses. We thank Dr. Lauren Penfield for the drawings in Figures 2, 3, and 6. We thank Dr. Xun (Austin) Ding, Jacob Hardwood, Kristin Mercier, Marc Anthony Pastor and Yijing Li for technical assistance. Thanks to all the colleagues in the D. Montell lab for advice on the project and comments on the manuscript. We thank the labs that generously shared reagents with us. Thanks to the *Drosophila* stock centers and DHSB for supplying fly stocks and antibodies.

## Author contributions

Conceptualization: XG, MM, WD, MN, DJM

Methodology: XG, MM, WD, MN, AYT, DM, ZC, CM, DJM

Investigation: XG, MM, WD, MN, AYT, DM, ZC

Visualization: XG, MM, WD, MN, AYT, ZC

Funding acquisition: DJM, CM

Project administration: DJM

Supervision: DJM, CM

Writing: XG, MM, WD, MN, ZC, DJM, CM

## Data and materials availability

All data are available in the main text or the supplementary materials.

## Supplementary Materials

Materials and Methods

Figs. S1 to S5

References (26–32)

**Supplementary Fig. 1.**
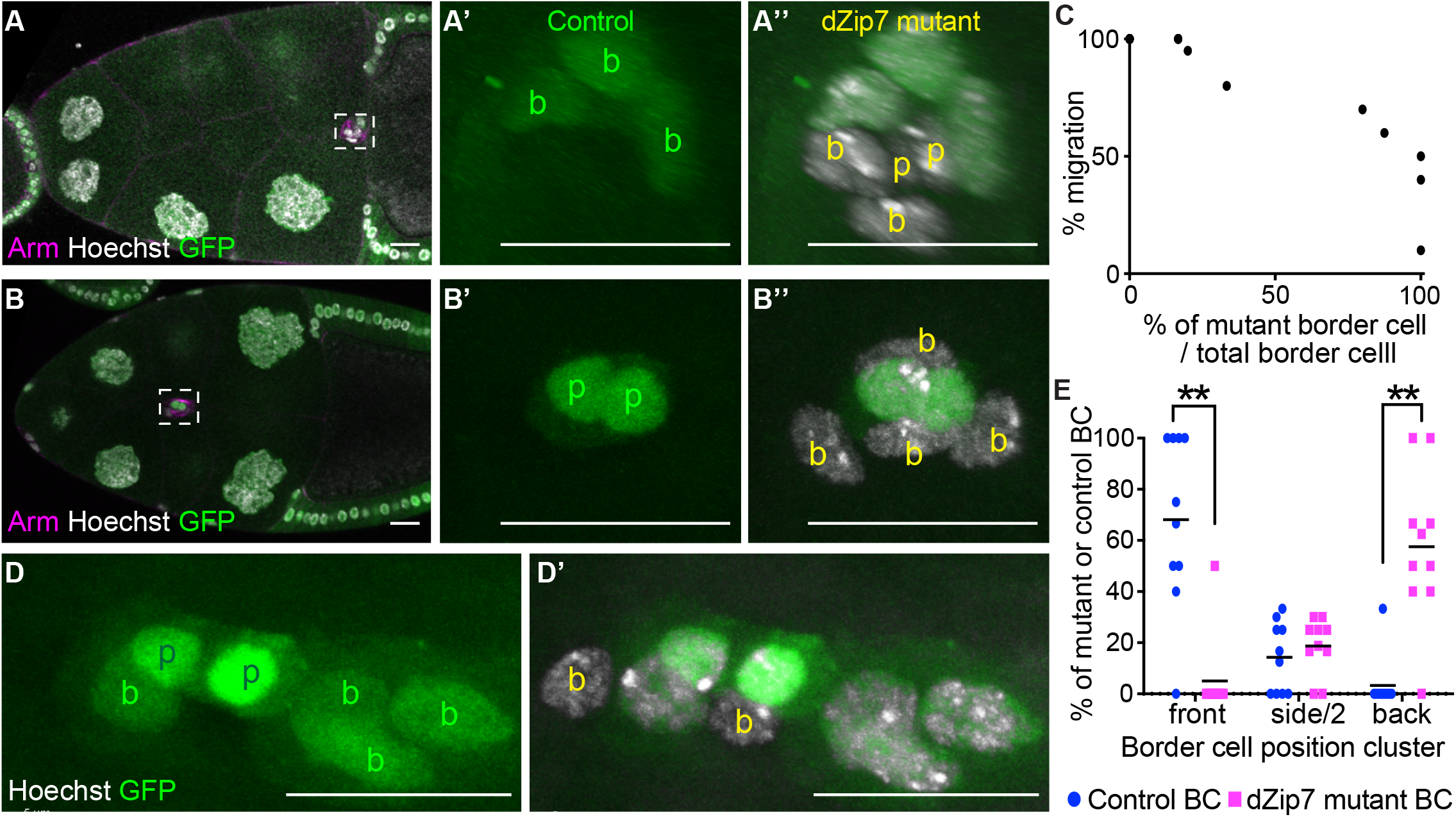
dZIP7 functions in migratory border cells (A-A”) An egg chamber with homozygous dZip7 mutant cells (GFP-). Both polar cells (p) and two border cells (b) are mutant. (B-B”) An egg chamber in which all outer border cells are GFP-/-(homozygous dZip7 mutant). (C) Migration distance expressed as a percentage of the migration path for mosaic border cell clusters as a function of the proportion of homozygous mutant cells in each cluster. (D) High magnification view showing the spatial distribution of dZip7+ (GFP+) and dZip7-/- (GFP-/-) cells in a migrating cluster. (E) Quantification of the percentage of dZip7+ vs dZip7-/-border cells in the front, side, or back of the border cell cluster showing that dZip7-/-cells are more likely to occupy a rear position. “p” indicated polar cells, “b” indicated border cells, green labels control cells, yellow labels mutant cells. ** P≤0.01, *** P≤0.001, **** P≤0.0001. Scale bars=20 μm.

**Supplementary Fig. 2.**
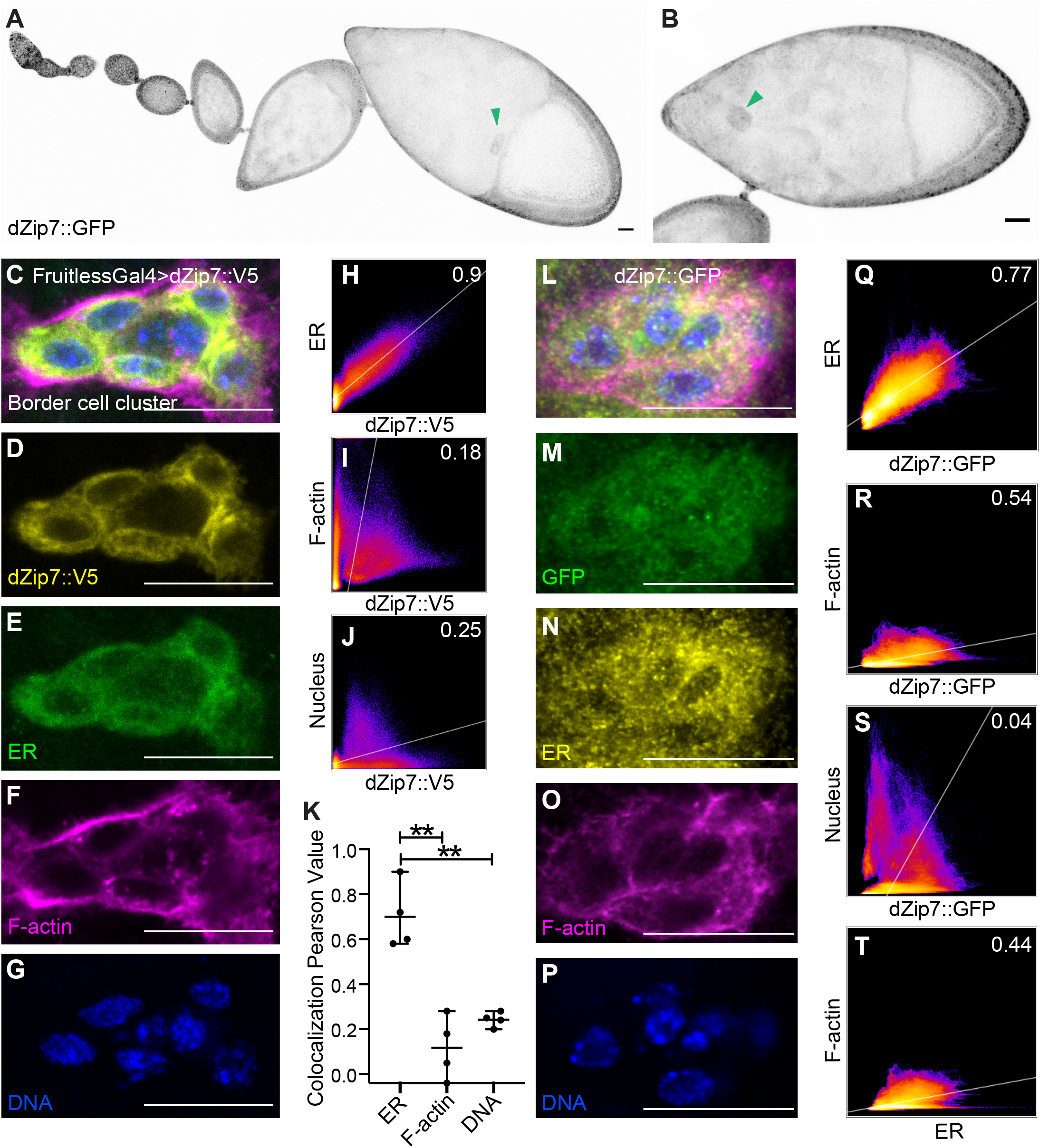
dZIP7 localizes to the ER (A, B) dZip7::GFP grayscale single channel corresponding to Figure 1A. (C-G) High magnification of a border cell cluster showing the localization of overexpressed dZip7::V5 (yellow), anti-PDI staining for ER (green), phalloidin (magenta), and Hoechst (blue). (H-J) 2-dimensional intensity histograms showing localization of dZip7::V5 relative to ER, F-actin, and DNA. The colocalization regression Pearson’s coefficient is displayed in the upper right corner. (K) Comparison of Pearson’s coefficients (average of 4 border cell clusters). ** P value < 0.01. (L-P) High magnification of a border cell cluster expressing dZip7::GFP under the control of endogenous regulatory sequences (green), ER (PDI, yellow), F-actin (phalloidin, magenta) and DNA (Hoechst, blue). (Q-T) 2-dimensional intensity histograms showing colocalization and Pearson’s coefficient for dZip7::GFP relative to ER, F-actin, nuclei, as well as ER relative to F-actin. Scale bars=20 μm.

**Supplementary Fig. 3.**
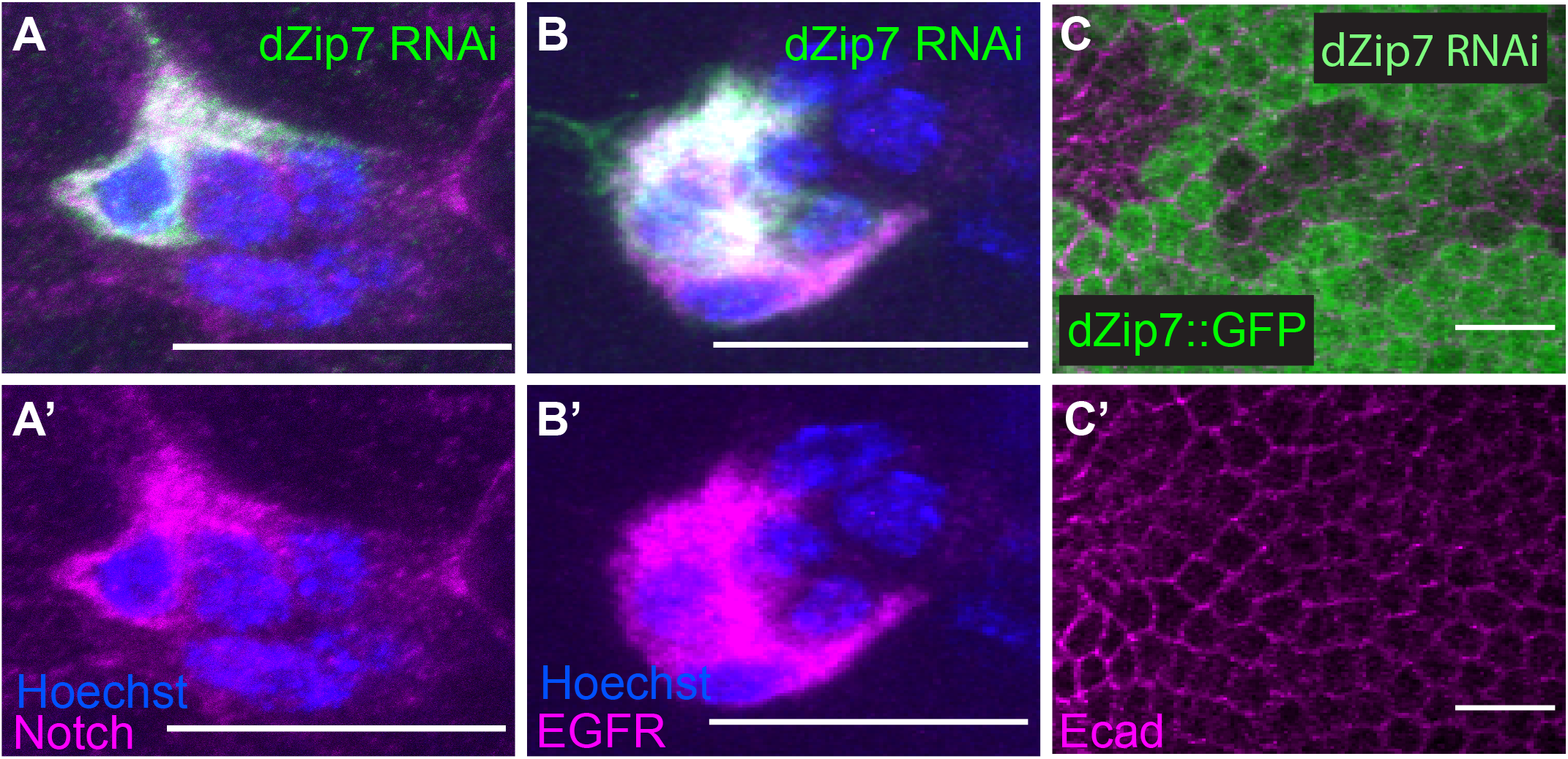
dZIP7 knockdown causes ER accumulation of Notch and EGFR but not Ecadherin. (A-A’) dZIP7RNAi-expressing clones (GFP+, green) accumulate intracellular Notch protein in border cells relative to neighboring wild type cells. (B, B’) Accumulation of EGFR (magenta) in dZIP7RNAi-expressing border cells. (C, C’) c306Gal4>dZip7RNAi reduces dZip7::GFP expression but does not cause E-cadherin (magenta) intracellular accumulation.Scale bars=20 μm.

**Supplementary Figure 4:**
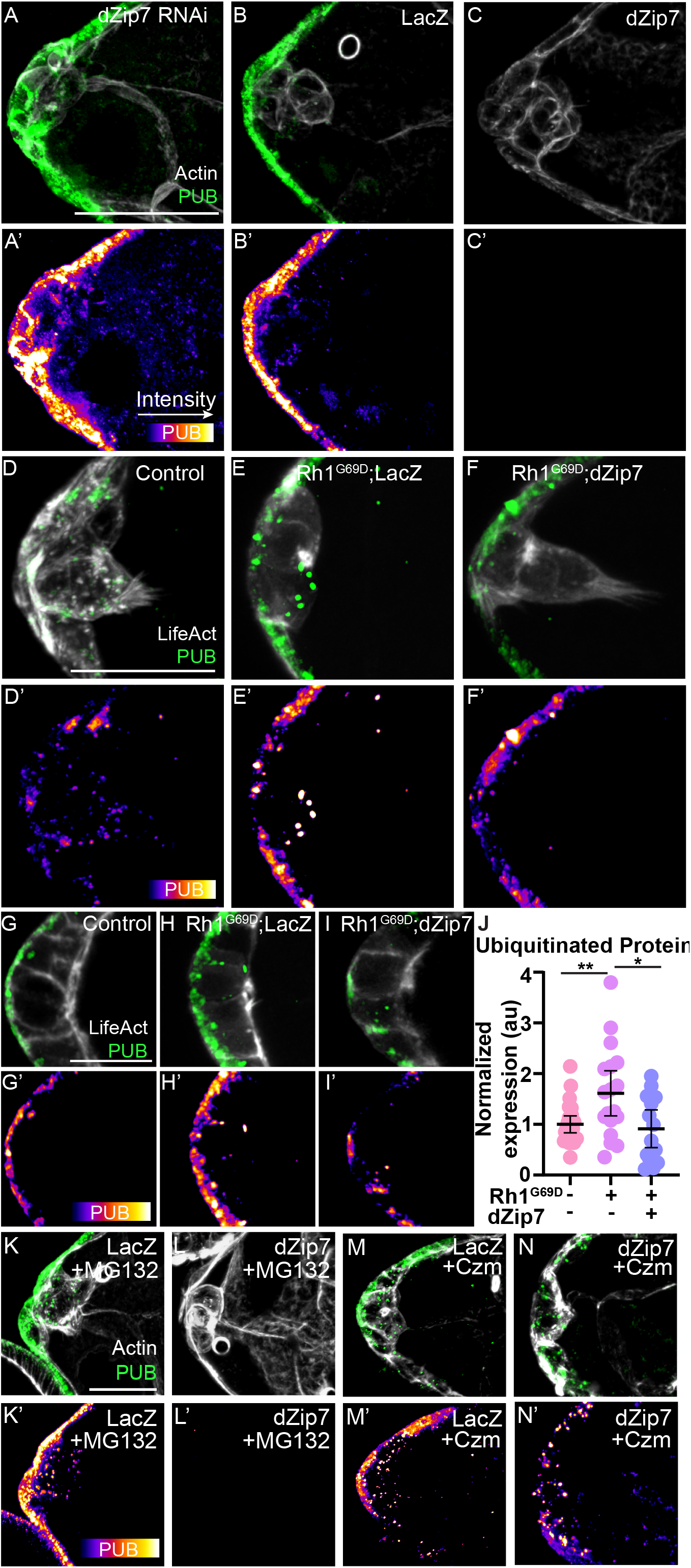
(A-C’) Representative images of stage 8 egg chambers stained with an antibody against ubiquitinated proteins (PUB). Scale bars, 10 μm. (D) Quantification of fluorescence intensity of ubiquitinated proteins. Expressing Rh1G69D causes buildup of ubiquitinated proteins compared to the control, but this effect is suppressed by co-expressing dZip7. *P≤0.05, ** P≤0.01. Error bars represent the 95% confidence intervals. Dots represent individual border cell clusters. (E-N) Representative images of stage 9 border cell clusters stained against ubiquitinated proteins. (E-G’) Co-expressing dZip7 and Rh1G69D reduced buildup of polyubiquitinated protein in border cells. (H-J’) dZip7 overexpression prevented buildup of ubiquitinated protein, while dZip7 knockdown increased ubiquitinated protein load. (K-L’) dZip7 overexpression also reduced ubiquitinated proteins in egg chambers treated with MG132. (M-N’) However, dZip7 overexpression does not prevent ubiquitinated protein buildup in egg chambers treated with Rpn11 inhibitor Capzimin (Czm). Scale bars=20 μm.

**Supplementary Figure 5:**
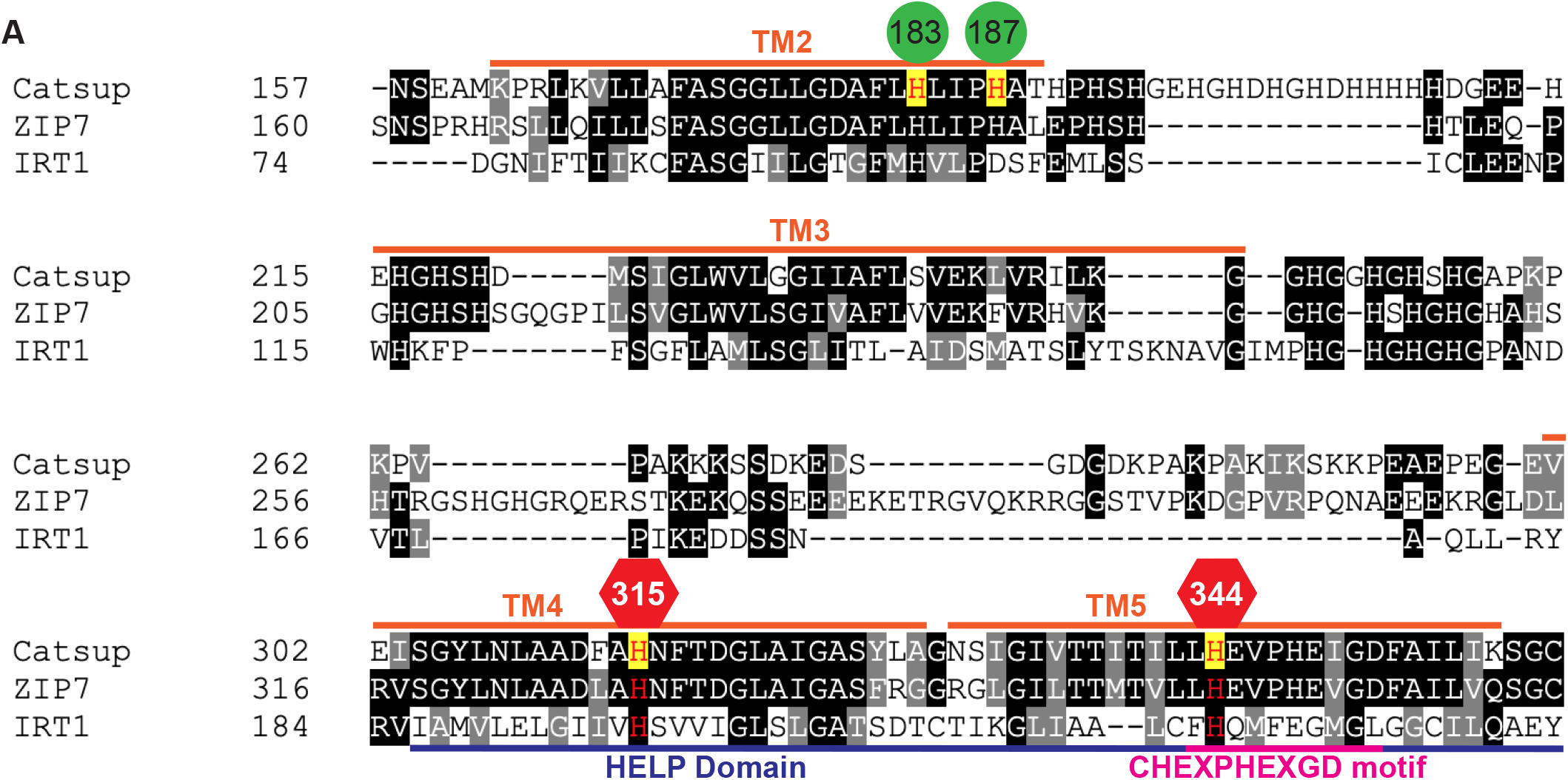
Histidine to alanine mutations predicted to disrupt Zn2+ transport (red) or not (green) indicated on the sequence alignment between plant IRT1, human ZIP7, and Drosophila dZIP7 (Catsup).

